# Pervasive impacts of climate change on the woodiness and ecological generalism of dry forest plant assemblages

**DOI:** 10.1101/2022.03.04.482968

**Authors:** Mario R. Moura, Fellipe A. O. do Nascimento, Lucas N. Paolucci, Daniel P. Silva, Bráulio A. Santos

## Abstract

Climate emergency is a significant threat to biodiversity in the 21^st^ century, but species will not be equally affected. In summing up different species’ responses at the local scale, we can assess changes in the species quantity and composition of biotic assemblages. Here we investigated climate change driven variation in species richness and spatial beta-diversity using modelled distributions of 2,841 plant species in Caatinga, the largest dry forest region of South America. More than 99% of plant assemblages were projected to lose species by 2060, with biotic homogenisation ─ the decrease in spatial beta-diversity forecasted in 40% of the Caatinga. Replacement of narrow-range woody species by wide-range non-woody ones should impact at least 85% of Caatinga plant assemblages. The future increase in aridity will change patterns of woodiness and ecological generalism of tropical dry forest plant assemblages, and ultimately erode ecosystem services linked to biomass productivity and carbon storage.

---

Climate change has been altered the environmental conditions experienced by many species on Earth ^1^. If species tolerances do not encompass the novel climatic conditions, they may be forced to change their phenology or geographic range to track suitable climates ^2,3^. Spatial changes in the geographic range of species can alter the composition of species assemblages ^4^. While certain species can colonise new sites in the future, most may not disperse quickly enough to avoid local extinctions, with the extinction risk being greatest for species with low vagility and narrow distribution ^5^. High local extinctions of narrow-range species and the potential colonisation of new sites by wide-range species can lead to the biotic homogenisation of species assemblages ^6^, and the eventual loss of ecosystem functions provided by such species ^7^. Because the climate emergency is a higher threat for tropical species ^2,5^, long-term conservation planning will benefit from understanding how different tropical ecosystems are subject to biotic changes ^8,9^.

Climate change has induced the biotic homogenisation of plant assemblages in several ecosystems around the world ^10^, including drylands ^8,11,12^. It has been suggested that dryland plants already experience a high water deficit and are close to their climatic tolerances ^9^. One of the world’s largest and floristically richest tropical dry forest is found in northeastern Brazil ─ the Caatinga ─ with 912,529 km^2^ ^13,14^. Future climate projections indicate increases in aridity, with subsequent desertification of some areas within the Caatinga ^15^. Previous research has shown that climate change should drive range contraction of endemic Caatinga plant species, especially those with more specialised life history attributes ^16^. Indeed, the colonisation and extinction rates of species assemblages may be affected by species geography and life-history attributes ^17^. Narrow-range species tend to be more sensitive to climate change, whereas wide-range species often exhibit broader climatic niches and thus high ecological generalism ^18^. Among flowering plants, the growth form is known to reflect species ecophysiology ^19^, with woody plant species likely exhibiting limited adaptability to climate change due to their longer generation time and slower rates of climatic niche evolution relative to non-woody plants ^20^.

We applied ecological niche models (ENMs) under an ensemble modelling framework to estimate the current and future geographic distribution patterns of 2,841 Caatinga plant species, and then assessed potential biotic changes in local plant assemblage richness (ΔS = S_future_ – S_current_) and spatial beta-diversity (Δβ_SOR_ = β_SOR.future_ – β_SOR.current_) in response to climate change (see Methods). Our investigation considered the latest projections on future climate scenarios ^21^ for 2060 and 2100, under the business-as-usual (SSP245) and non-mitigation (SSP585) scenarios. Ensemble models showed good predictive performance, with an average Sørensen similarity index of 0.934 (SD = 0.043, range = 0.703–1.00; Fig. S2). Because our results were qualitatively similar for 2060 and 2100, we focused on 2060 projections for brevity (see Extended Data for results concerning the year 2100).

## RESULTS

Climate change will drastically alter plant biodiversity in one of the world’s largest seasonal tropical dry forests, the Caatinga. Our projections show that almost 90% of Caatinga plant species will lose suitable areas by 2060, particularly narrow-range species (Fig 1). The current distribution of Caatinga plant species will decrease on average by 37.4% and 43.9% in the SSP245 and SSP585 scenarios, respectively. No Caatinga plant species is projected to lose its entire suitable area within the Neotropics, but in the Caatinga, from 62 (SSP245) to 89 (SSP585) species could be regionally extinct ─100% of range loss─ by 2060, and between 141 (SSP245) and 349 (SSP585) species could be regionally extinct by 2100.

**Fig 1.**
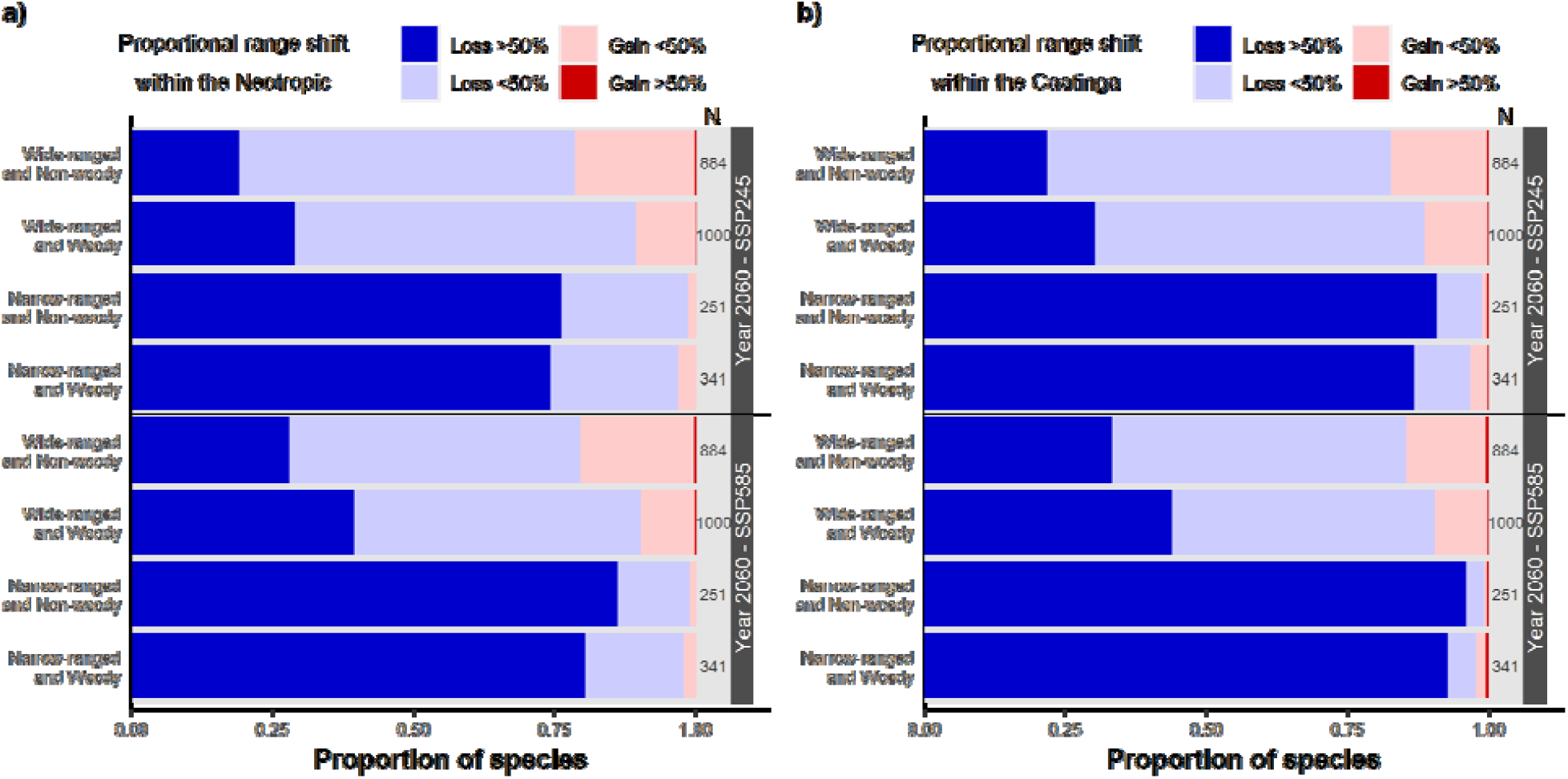
Projected range shift for species holding different levels of woodiness and ecological generalism. Range shifts were computed separately within the (a) Neotropics and (b) Caatinga. The results are shown for 2060 under the business-as-usual (SSP245) and non-mitigation (SSP585) scenarios. See Fig. S3 for results concerning 2100.

At the assemblage-level, more than 99% of plant assemblages in the Caatinga will lose species by 2060 (Fig. 2b and c). Biotic homogenisation (Δβ_SOR_ < 0) is expected in about 40% of Caatinga plant assemblages, particularly in species-poor regions currently dominated by non-woody and wide-range species (Fig. 2e and f). Relative to regions subject to biotic heterogenisation, the future homogenised plant assemblages currently harbour lower species richness (χ^2^ = 3834.9, df = 7, p < 0.001, Fig. 3a), lower proportion of woody species (χ ^2^ = 1008.6, df = 7, p < 0.001, Fig. 3b), and higher proportion of wide-range species (χ ^2^ = 1953.1, df = 7, p < 0.001, Fig. 3c). Although we projected a pervasive decrease in woodiness and an increase in the ecological generalism of plant assemblages, the magnitude of such changes differs between future homogenised or heterogenised regions. Plant assemblages facing homogenisation risk by 2060 showed lower species loss (χ^2^ = 4478.8, df = 7, p < 0.001, Fig. 3d), higher decrease in relative contribution of woody species (χ ^2^ = 1419.3, df = 7, p < 0.001, Fig. 3e), and lower increase in relative contribution of wide-range species (χ ^2^ = 1662.6, df = 7, p < 0.001, Fig. 3f) than assemblages subject to heterogenisation.

**Fig 2.**
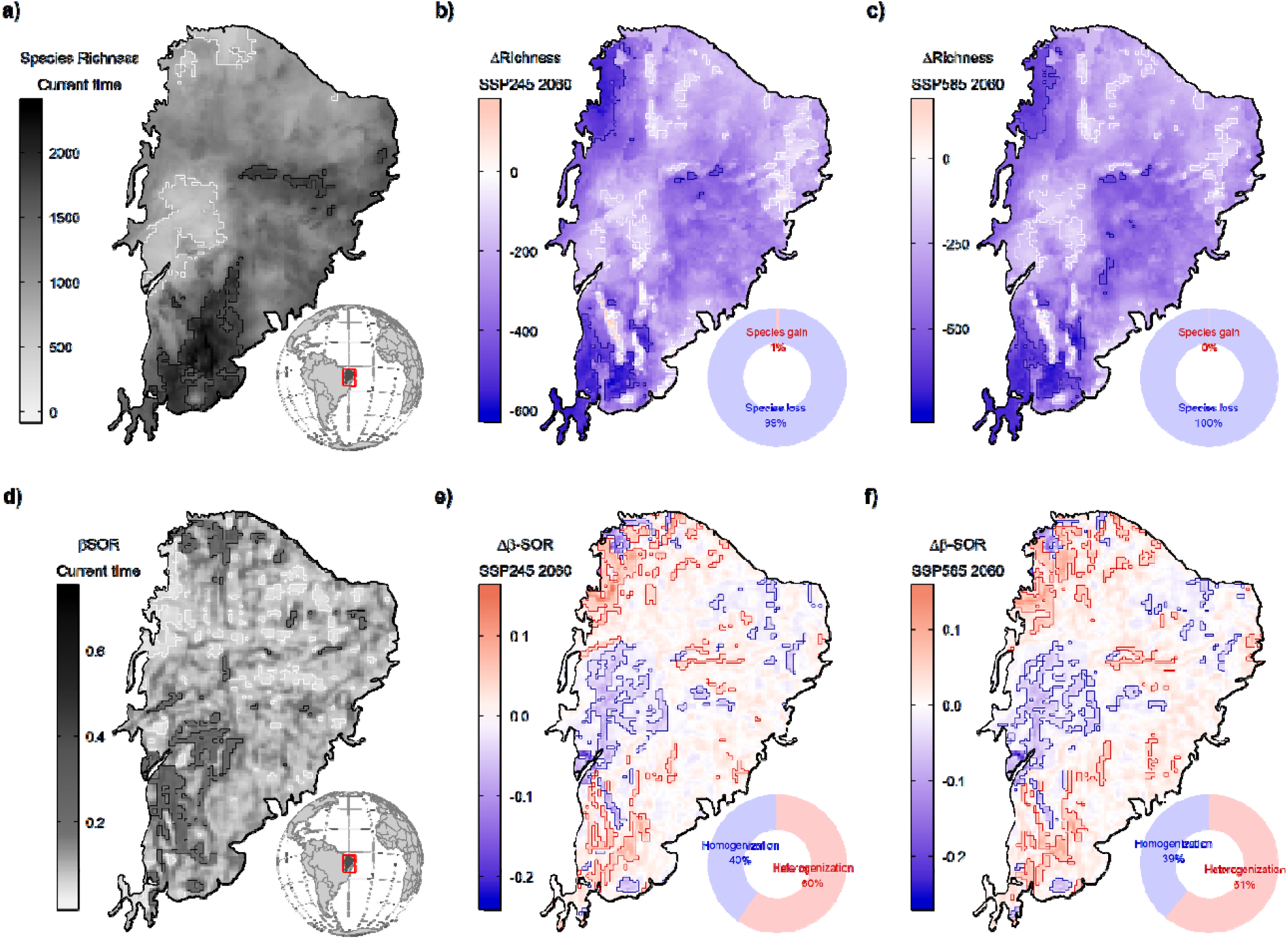
Geographical patterns of plant species richness and spatial beta-diversity in the Caatinga. (a) Projected species richness at the current time. Expected change in species richness (ΔS) across plant assemblages under the (b) business-as-usual (SSP245) and (c) non-mitigation (SSP585) scenarios in 2060. (d) Spatial beta-diversity (β_SOR_) for the current time. Expected change in spatial beta-diversity (Δβ_SOR_) across plant assemblages under (e) SSP245 and (f) SSP585 scenarios for 2060. The contour lines denote the assemblages (cells) in the upper and lower 10% of the mapped pattern. See Fig. S4 for results concerning 2100.

**Fig 3.**
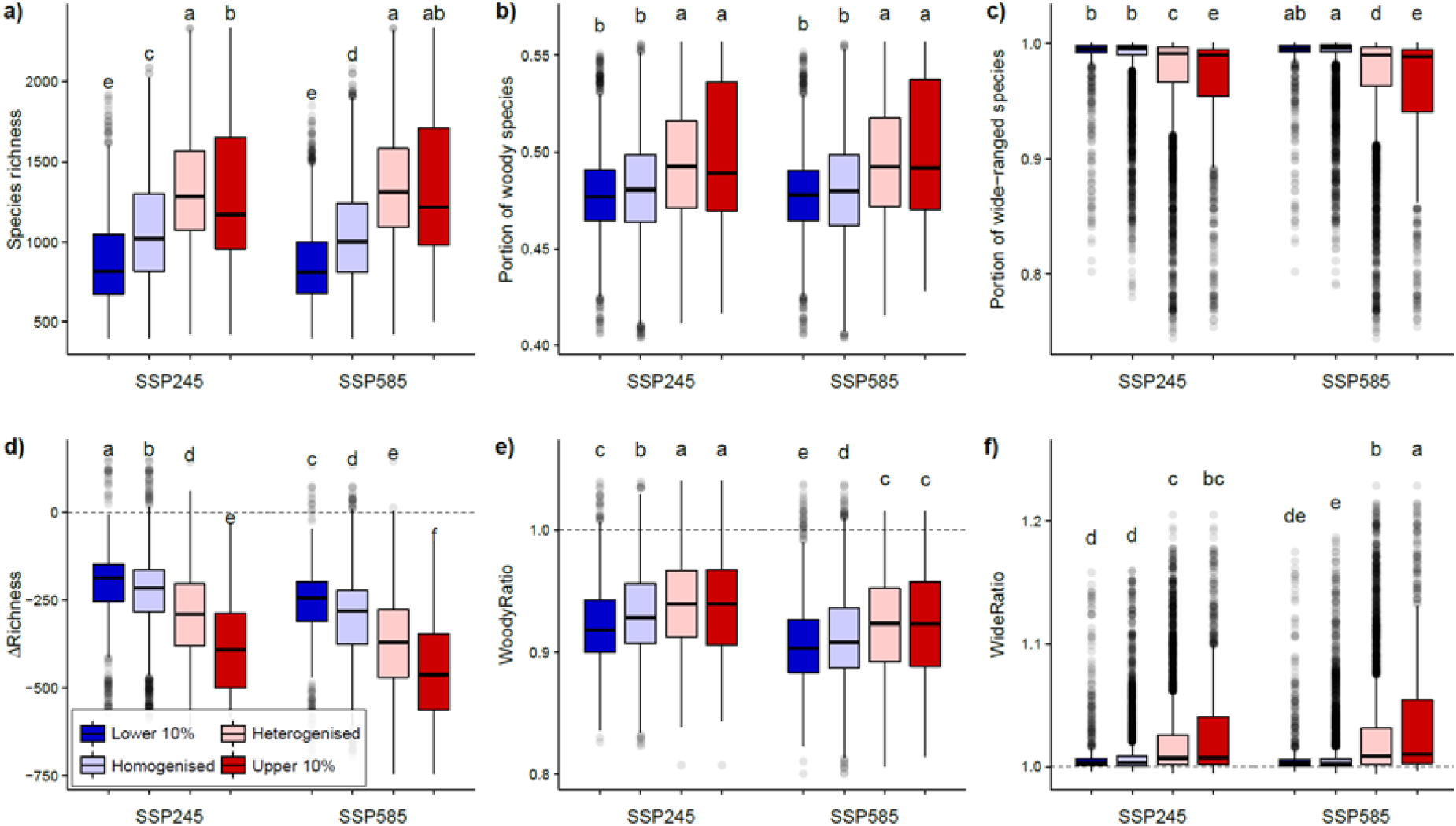
Assemblage-level metrics across regions subject to different levels of projected biotic change by 2060. Each box denotes the median (horizontal line) and the 25th and 75th percentiles. Vertical lines represent the 95% confidence intervals, and black dots are outliers. Small capital letters denote the results of the Kruskal–Wallis tests for the difference in medians across assemblages subject to different levels of biotic homogenisation (p = 0.05, using Bonferroni correction). Woody and wide ratios above 1 indicate an increase in the assemblage-level proportion of woody and wide-range species in the future. See Fig. S5 for results concerning 2100.

We observed a predominance of assemblages with a higher proportion of non-woody and wide-range species in the northern and middle-west regions of the Caatinga, whereas assemblages with relatively more woody and narrow-range species occurred in the southern and northeastern Caatinga (Fig. 4). Under the business-as-usual scenario, 98.4% of plant assemblages will experience a reduction in the proportion of woody species (WoodyRatio < 1), slightly less than in the non-mitigation scenario (98.9% of assemblages). The relative contribution of wide-range species will increase (WideRatio > 1) in most assemblages in both SSP scenarios (86.1% in the SSP245 and 85.3% in the SSP585, Fig. 4b and c). In all SSP scenarios investigated, the increase in spatial beta-diversity of plant assemblages was directly related to species loss (Fig. 5a and b), with changes in relative contribution of wide-range species linked to the decrease in proportion of woody species (Fig. 5c and d).

**Fig 4.**
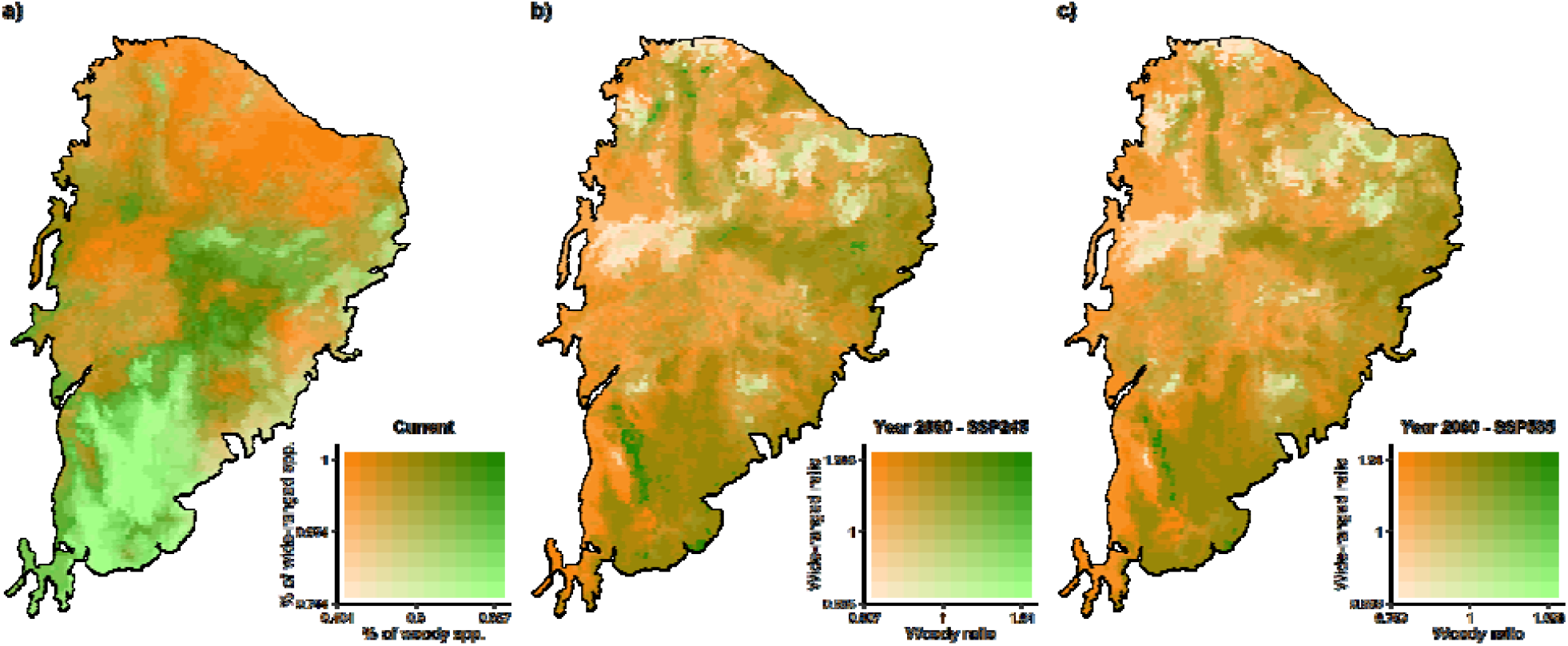
Patterns of woodiness and ecological generalism of plant assemblages in the Caatinga. (a) Proportion of woody and wide-ranged species in plant assemblages. Relative change in the proportion of woody and wide-range species between 2060 and the current time under the (b) business-as-usual, SSP245 and (c) non-mitigation, SSP585 scenarios. Woody and Wide-range ratios above 1 indicate an increase in the assemblage-level proportion of woody and wide-range species in the future. See Fig. S6 for results concerning 2100.

**Fig 5.**
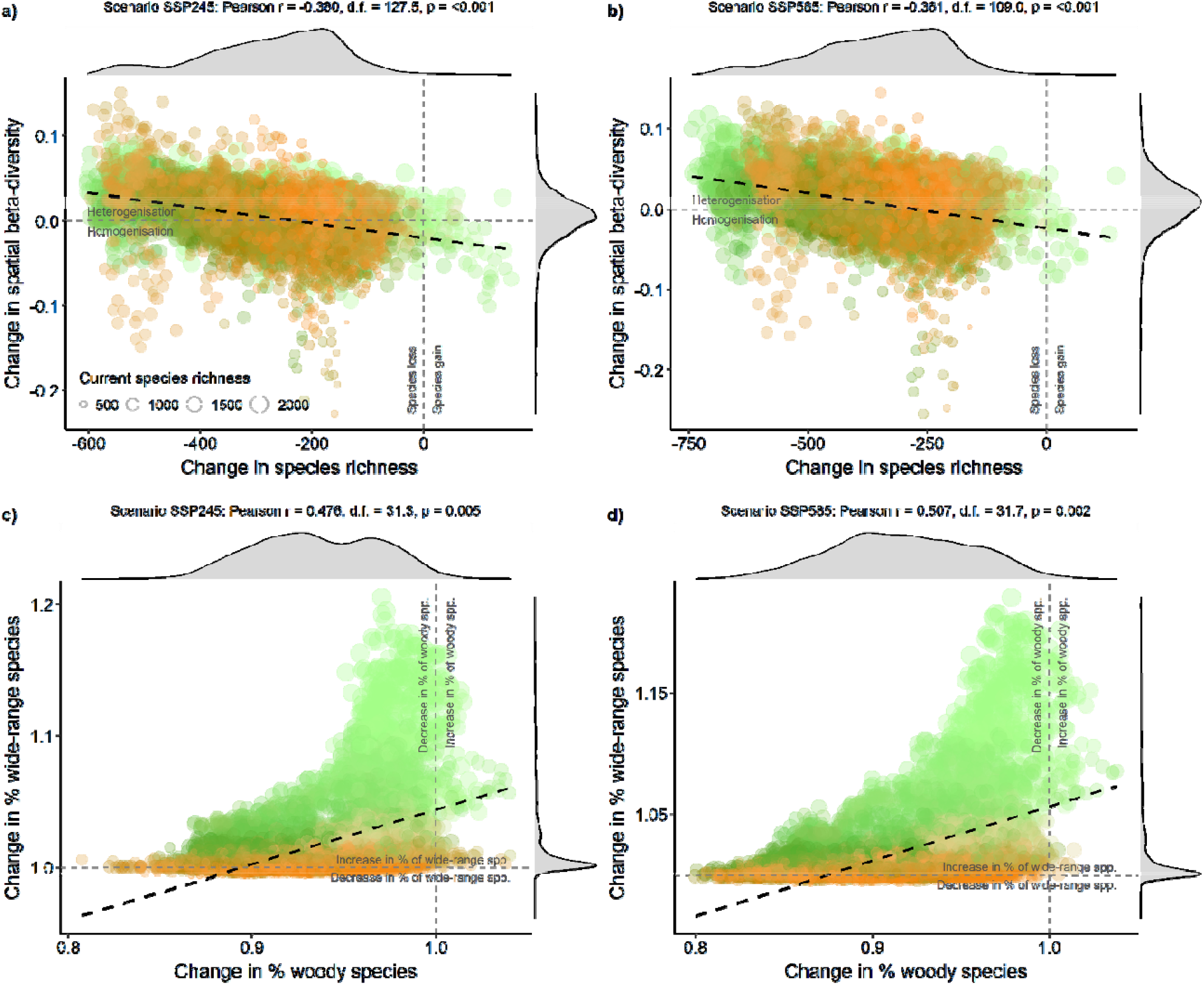
Projected change in species richness, spatial beta-diversity, woodiness, and ecological generalism in Caatinga plant assemblages by 2060. Relationship between differences in species richness (ΔS) and spatial beta-diversity (Δβ_SOR_) in the scenarios (a) Business-as-usual, SSP245, and (b) Non-mitigation SSP585. Relationship between change in relative contribution of woody (WoodRatio) and wide-range (WideRatio) species. Symbol colours follow species assemblage representation in Fig. 4a. Pearson correlations on the top of each panel were based on spatially corrected degrees of freedom. See Fig. S7 for results concerning 2100.

## DISCUSSION

The exacerbated decrease in plant richness can erode ecosystem services in Caatinga by 2060. In drylands worldwide, the role of plant richness in productivity stability is as important as that of climate and edaphic conditions ^22^. Under high aridity, species-rich assemblages are more critical for ecosystem stability, whereas functionally distinct species minimise variation in the temporal delivery of ecosystem services at low aridity ^22^. Climate change is expected to increase aridity in Caatinga, particularly in the central-southern region ^15^, where our projections indicate a higher species loss of plant assemblages. To worsen the situation, 98.4–98.9% of Caatinga plant assemblages will lose relatively more woody than non-woody species, which should enhance the impacts on biomass productivity and carbon storage in drylands ^23,24^.

Aridity may favour the establishment of wide-range plant species ^25^. Since most wide-range plant species in Caatinga have non-woody growth-forms ^26^, the projected increase in aridity in this region will lead to structural changes in vegetation complexity. With higher aridity, dryland ecosystems face a vegetation decline phase due to the reduction of leaf area and canopy cover ^27^. Aridification can also promote compositional change ^28^ and reduce the beta-diversity of dryland plant assemblages ^29^. As environmental filtering can better explain the beta-diversity of herbs and shrubs than that of trees ^30,31^, woody species’ distributions are likely at a lower equilibrium with climate than those of non-woody species, implying that woody species may not keep pace with climate change. Projected changes in the species richness and beta-diversity of Caatinga plant assemblages can therefore underestimate the impacts of climate change on plant assemblages with higher levels of woodiness.

The impacts of climate change are often expected to be less severe in mountainous regions ^32^. Although elevational gradients can allow species to track more suitable climates over time, the spatial configuration of mountainous areas can limit elevational range shifts ^33^, particularly for woody species ^34^. In the Caatinga, the four most relevant highlands ─ where many narrow-range woody species concentrates ─ are disconnected from each other and located in transitional zones in the south (*Chapada Diamantina*), east (*Planalto da Borborema*) and central-northwest (*Chapada do Araripe* and *Serra da Ibiapaba*), which impose additional dispersal constraints on woody species there. Threats to woody species and their ecosystem functions could be considerably greater than those we project. Vertebrates and invertebrates interacting with woody plants ─ herbivores, seed dispersers, and pollinators ─ will have to couple with the sudden changes in availability and composition of plant resources ^35,36^, ultimately scaling up the potential for disruption of biotic interactions ^37^.

We have shown how climate change can jeopardise Caatinga plant biodiversity, but much of this region is also affected by chronic disturbances ^38^ that can operate synergistically with climate change, and intensify the impacts of biodiversity loss on ecosystem functions ^39^. Caatinga already lost half its original cover, and more than 90% of the remaining fragments have less than 500 ha. ^38^. For some non-woody, self-pollinated, and wind-dispersed species, this scenario may not prevent range expansion, but our projections show an extensive reduction in suitable areas for most woody and non-woody species. Interestingly, the western half of the Caatinga, where biotic changes are most pronounced, also concentrates the most conserved dry forest remnants ^38^. This coincident spatial configuration imposes both a challenge and an opportunity to expand the protected area network to assure the connectivity and long-term persistence of Caatinga plant assemblages. Caatinga still figures with around 1% of the original extension covered by strictly protected areas ^40^, far beyond the more recent thresholds established under the post-2020 Global Biodiversity Frame of conserving 30% of Earth’s land by 2030 ^41^. As a member of the Convention on Biological Diversity and the sole holder of the Caatinga, Brazil will have a crucial role in conserving the most extensive tropical dry forest in South America.

## METHODS

### Species data

We compiled occurrence records of Caatinga flowering plants from the scientific literature and herbarium records, available at the Global Biodiversity Information Facility ^42^ and speciesLink (splink.cria.org.br). We restricted the spatial coverage of the species occurrence dataset to the Neotropical region and recorded a total of 4,890,681 occurrences for 8,629 species. Records that were duplicated, with georeferencing errors or uncertain identification were excluded. To reduce the potential effect of sampling bias and spatial autocorrelation in the occurrence dataset, we randomly filtered one occurrence record for each species within a radius of ∼10 km ^43^ leading to 1,024,363 occurrences of 7,936 species. Preliminary inspections indicated that many species occurred marginally in the Caatinga. Because our focus was on typical Caatinga plants, we kept only those species with at least 10% of their occurrences within the Caatinga, resulting in 345,848 occurrences of 4,534 species. We excluded species with fewer than 15 occurrence records ^44^ and kept for subsequent modelling procedures 335,091 records of 2,841 species belonging to 776 genera and 141 botanical families (Fig. S1).

We gathered data on the growth-form of Caatinga flowering species using the Botanical Information and Ecology Network ^45^, the Plant Trait Database ^46^, and the Brazilian Flora 2020 ^47^, complemented by pertinent literature ^26,48,49^. For each species, we assigned one out of seven growth form types: tree, shrub, palm tree, woody vine, herb, herbaceous vine, or succulent. For species with multiple growth-form categories we assigned the growth form agreed upon by most sources ^50^. We then grouped the species into two categories: (i) woody (trees, shrubs, palms, and woody vines) and (ii) non-woody species (herbs, herbaceous vines, and succulents). Growth-form was assigned to 2,476 angiosperm species (1,341 woody and 1,135 non-woody species, Table S1), covering 87.2% of species in our database.

### Current and future climate projections

We used the 19 bioclimatic variables from the WorldClim database, version 2.1 ^51^ to represent the current climate. Bioclimatic layers were downloaded at 5 arc-min (∼10 km) and cropped to the extent of the Neotropical realm (our background). To avoid problems with multicollinearity and reduce the dimensionality of predictor layers, we performed a principal component analysis (PCA) and retained the first six axes representing the principal components as climate predictors. PCA axes explained 96.8% of the variation in the original data. We used PCA loading coefficients to project the linear relationship between raw predictors and principal components onto new layers representing future climate scenarios.

We considered future climate projections for the periods 2041–2060 (hereafter 2060) and 2081–2100 (hereafter 2100) following the 6th Assessment Report of the Intergovernmental Panel on Climate Change ^21^. We used two Shared Socioeconomic Pathways (SSP) for each period: SSP245 and SSP585 as a business-as-usual and non-mitigation scenario, respectively. Because the selection of different generalised circulation models (GCM) is recognised as a source of uncertainty in projecting the future habitat suitability of species ^52^, we used five GCMs for each combination of the period and SSP, namely: BCC-CSM2-MR, CNRM-CM6-1, IPSL-CM6A-LR, MIROC6, and MRI-ESM2-0.

### Ecological niche models

For each species, we computed pseudoabsences using the same number of observed presences to maintain the presence-absence ratio of 1:1 ^53^. Pseudoabsences were allocated following the environmentally constrained method, based on the lowest suitable region predicted by a climate envelope ^54^. The choice of the statistical method or algorithm can affect the resulting predictions from an ecological niche model (ENM) depending on the initial modelling conditions ^55^. We computed an ensemble of projections for each species to minimise uncertainty around the ENM method ^56^. The ensemble included projections with six methods: Climate envelope (BIOCLIM), Gower Environmental Distance (DOMAIN), Generalised Linear Models, Generalised Additive Models, Maximum Entropy, and Random Forests. We used the species accessible area to mask its respective projections to avoid predicting habitat suitability for regions unreachable by a species within the time frame of projected climate change. The accessible area for each species was defined by a buffer with a width size equal to the maximum nearest neighbour distance among pairs of occurrences ^57^. Models were calibrated for the baseline period using 4-fold cross-validation, 75% of randomly selected samples used for model training and the remaining 25% used for testing in each iteration.

To evaluate model performance, we measured the similarity between predictions and observations using the Sørensen similarity index, which is independent of species prevalence ^58^. It is necessary to binarise the species habitat suitability according to some threshold value to compute the Sørensen index. We used the species suitability value that maximised the Sørensen index for each algorithm at the baseline period. The ensemble model for each species was computed as the average weighted suitability, with weights given by the Sørensen index calculated for each algorithm. We used the average binarisation threshold weighted by the Sørensen index to binarise the ensemble habitat suitability into presence-absence maps for each species in current time and future scenarios.

We applied an occurrence-based restriction to keep only the patches of suitable habitat considered reachable by a species ^59^ to minimise overprediction issues associated with presence-absence maps derived from ENMs. Patches of suitable habitats are assumed to be reachable by the species if they overlap with a presence record or are within an edge-edge distance threshold of an occupied suitable patch ^59^. This distance threshold was defined as the maximum nearest neighbour distance among pairs of occurrences of the respective species. Computations were performed in R 4.1.0 ^60^ using the package *ENMTML* ^61^.

### Spatial patterns of beta-diversity, woodiness, and ecological generalism

We mapped the Caatinga using an equal-area projection grid cell of 10×10 km of spatial resolution to assess changes in beta-diversity spatial pattern. Using the binary maps for the ensemble projection, we built species presence-absence matrices for projections based on the current time and for each combination of the future period (2060 and 2100) and the SSP scenario (SSP245 and SSP585). We only considered species presence in a grid cell if they occupied at least 50% of the cell area. The spatial beta-diversity for each grid cell was measured by the Sørensen-based multiple-site dissimilarity index, β_SOR_ ^62^, computed for the cell set formed by the focal cell and its immediately adjacent neighbour cells. Because the size of the cell set can affect the β_SOR_ value ^63^, we applied a subsampling procedure to randomly select four neighbour cells around each focal cell 100 times to compute the average β_SOR_ across iterations. We used the β_SOR_ difference between each future and current scenario (Δβ_SOR_ = β_SOR.future_ – β_SOR.current_) to identify plant assemblages (cells) subject to biotic homogenisation (Δβ_SOR_ < 0) or heterogenisation (Δβ_SOR_ > 0). We also calculated the difference between future and current species richness (ΔS = S_future_ – S_current_). Computations were performed in R using the *betapart* package ^64^.

We used the species growth form to compute the proportion of woody species in each plant assemblage (WoodyProp) to assess potential changes in the assemblage-level patterns of woodiness and ecological generalism. To measure the assemblage-level ecological generalism, we initially classified as narrow-range plant species whose range size distribution was below the first quartile (∼100 mil km^2^) of current projected distribution within Caatinga, or otherwise wide-range. Then, we extracted the proportion of wide-range species in each plant assemblage (WideProp). We calculated WoodyProp and WideProp for the current and future scenarios and used the ratio of future to current time to represent the relative change in woodiness (WoodyRatio = WoodyProp_future_ / WoodyProp_current_) and ecological generalism (WideRatio = WideProp_future_ / WideProp_current_) of plant assemblages. WoodyRatio and WideRatio above 1 indicate a future increase in the assemblage-level proportions of woody and wide-range species, respectively. We used Kruskal-Wallis tests to assess whether the medians of (i) Current species richness, (ii) ΔS, (iii) WoodyProp_current_, (iv) WoodyRatio, (v) WideProp_current_, and (vi) WideRatio differ between assemblages subject to biotic homogenisation (Δβ_SOR_ < 0) or heterogenisation (Δβ_SOR_ > 0). Linear relationships between changes in species richness (ΔS) and spatial beta-diversity (Δβ_SOR_), and changes in the relative contribution of woody (WoodyRatio) and wide-range (WideRatio) species were verified through a modified t-test ^65^ to spatially correct the degrees of freedom of correlation coefficients. Computations were performed in R using the packages *SpatialPack* ^66^.

## ACKNOWLEDGMENTS

We are grateful to E. G. Moura-Jr for suggestions in previous versions of the manuscript. To Coordenação de Aperfeiçoamento de Pessoal de Nível Superior (CAPES) for fellowships made available to FAON. To Conselho Nacional de Desenvolvimento Científico e Tecnológico (CNPq) for research grants in support of BAS (#312178/2019-0) and DPS (#304494/2019–4). To Universidade Federal da Paraíba for research grants in support of BAS (PVA-13357-2020).

## AUTHOR CONTRIBUTIONS

MRM, FAON, and BAS conceived the study; FAON compiled the data, MRM and FAON analysed the data. MRM developed the figures and led the writing. All authors contributed critically to the drafts and gave final approval for publication.

## ADDITIONAL INFORMATION

Supplementary Information is available for this paper, including Supplementary Tables (S1–S2) and Supplementary Figures (S1–S7).

