## Supplementary Information for "Pervasive impacts of climate change on the woodiness and ecological generalism of dry forest plant assemblages"

**This PDF file includes:**

Supplementary Tables

Supplementary Figures

### SUPPLEMENTARY TABLES

**Table S1. Proportion of Caatinga plant species showing projected range shift across different categories of woodiness and ecological generalism.** SSP denotes the future scenarios of Shared Socioeconomic Pathways. Range shift calculations considered the Neotropical extent.

| Woodiness | Generalism | Range shift Category | Number of Cells | % Species | Year | SSP |
|---|---|---|---|---|---|---|
| Non-woody | Narrow-ranged | Loss >50% | 251 | 0.908 | 2060 | ssp245 |
| Non-woody | Narrow-ranged | Loss <50% | 251 | 0.080 | 2060 | ssp245 |
| Non-woody | Narrow-ranged | Gain <50% | 251 | 0.008 | 2060 | ssp245 |
| Non-woody | Narrow-ranged | Gain >50% | 251 | 0.004 | 2060 | ssp245 |
| Non-woody | Wide-ranged | Loss >50% | 884 | 0.218 | 2060 | ssp245 |
| Non-woody | Wide-ranged | Loss <50% | 884 | 0.609 | 2060 | ssp245 |
| Non-woody | Wide-ranged | Gain <50% | 884 | 0.170 | 2060 | ssp245 |
| Non-woody | Wide-ranged | Gain >50% | 884 | 0.003 | 2060 | ssp245 |
| Woody | Narrow-ranged | Loss >50% | 341 | 0.868 | 2060 | ssp245 |
| Woody | Narrow-ranged | Loss <50% | 341 | 0.100 | 2060 | ssp245 |
| Woody | Narrow-ranged | Gain <50% | 341 | 0.029 | 2060 | ssp245 |
| Woody | Narrow-ranged | Gain >50% | 341 | 0.003 | 2060 | ssp245 |
| Woody | Wide-ranged | Loss >50% | 1000 | 0.303 | 2060 | ssp245 |
| Woody | Wide-ranged | Loss <50% | 1000 | 0.583 | 2060 | ssp245 |
| Woody | Wide-ranged | Gain <50% | 1000 | 0.111 | 2060 | ssp245 |
| Woody | Wide-ranged | Gain >50% | 1000 | 0.003 | 2060 | ssp245 |
| Non-woody | Narrow-ranged | Loss >50% | 251 | 0.960 | 2060 | ssp585 |
| Non-woody | Narrow-ranged | Loss <50% | 251 | 0.032 | 2060 | ssp585 |
| Non-woody | Narrow-ranged | Gain <50% | 251 | 0.004 | 2060 | ssp585 |
| Non-woody | Narrow-ranged | Gain >50% | 251 | 0.004 | 2060 | ssp585 |
| Non-woody | Wide-ranged | Loss >50% | 884 | 0.334 | 2060 | ssp585 |
| Non-woody | Wide-ranged | Loss <50% | 884 | 0.520 | 2060 | ssp585 |
| Non-woody | Wide-ranged | Gain <50% | 884 | 0.140 | 2060 | ssp585 |
| Non-woody | Wide-ranged | Gain >50% | 884 | 0.006 | 2060 | ssp585 |
| Woody | Narrow-ranged | Loss >50% | 341 | 0.927 | 2060 | ssp585 |
| Woody | Narrow-ranged | Loss <50% | 341 | 0.050 | 2060 | ssp585 |
| Woody | Narrow-ranged | Gain <50% | 341 | 0.018 | 2060 | ssp585 |
| Woody | Narrow-ranged | Gain >50% | 341 | 0.006 | 2060 | ssp585 |
| Woody | Wide-ranged | Loss >50% | 1000 | 0.440 | 2060 | ssp585 |
| Woody | Wide-ranged | Loss <50% | 1000 | 0.465 | 2060 | ssp585 |
| Woody | Wide-ranged | Gain <50% | 1000 | 0.092 | 2060 | ssp585 |
| Woody | Wide-ranged | Gain >50% | 1000 | 0.003 | 2060 | ssp585 |
| Non-woody | Narrow-ranged | Loss >50% | 251 | 0.976 | 2100 | ssp245 |
| Non-woody | Narrow-ranged | Loss <50% | 251 | 0.016 | 2100 | ssp245 |
| Non-woody | Narrow-ranged | Gain <50% | 251 | 0.004 | 2100 | ssp245 |
| Non-woody | Narrow-ranged | Gain >50% | 251 | 0.004 | 2100 | ssp245 |
| Non-woody | Wide-ranged | Loss >50% | 884 | 0.379 | 2100 | ssp245 |
| Non-woody | Wide-ranged | Loss <50% | 884 | 0.486 | 2100 | ssp245 |
| Non-woody | Wide-ranged | Gain <50% | 884 | 0.127 | 2100 | ssp245 |
| Non-woody | Wide-ranged | Gain >50% | 884 | 0.008 | 2100 | ssp245 |
| Woody | Narrow-ranged | Loss >50% | 341 | 0.950 | 2100 | ssp245 |

| | | | | | | |
|---|---|---|---|---|---|---|
| Woody | Narrow-ranged | Loss <50% | 341 | 0.032 | 2100 | ssp245 |
| Woody | Narrow-ranged | Gain <50% | 341 | 0.015 | 2100 | ssp245 |
| Woody | Narrow-ranged | Gain >50% | 341 | 0.003 | 2100 | ssp245 |
| Woody | Wide-ranged | Loss >50% | 1000 | 0.518 | 2100 | ssp245 |
| Woody | Wide-ranged | Loss <50% | 1000 | 0.399 | 2100 | ssp245 |
| Woody | Wide-ranged | Gain <50% | 1000 | 0.080 | 2100 | ssp245 |
| Woody | Wide-ranged | Gain >50% | 1000 | 0.003 | 2100 | ssp245 |
| Non-woody | Narrow-ranged | Loss >50% | 251 | 0.996 | 2100 | ssp585 |
| Non-woody | Narrow-ranged | Gain <50% | 251 | 0.004 | 2100 | ssp585 |
| Non-woody | Wide-ranged | Loss >50% | 884 | 0.630 | 2100 | ssp585 |
| Non-woody | Wide-ranged | Loss <50% | 884 | 0.282 | 2100 | ssp585 |
| Non-woody | Wide-ranged | Gain <50% | 884 | 0.072 | 2100 | ssp585 |
| Non-woody | Wide-ranged | Gain >50% | 884 | 0.016 | 2100 | ssp585 |
| Woody | Narrow-ranged | Loss >50% | 341 | 0.974 | 2100 | ssp585 |
| Woody | Narrow-ranged | Loss <50% | 341 | 0.015 | 2100 | ssp585 |
| Woody | Narrow-ranged | Gain <50% | 341 | 0.009 | 2100 | ssp585 |
| Woody | Narrow-ranged | Gain >50% | 341 | 0.003 | 2100 | ssp585 |
| Woody | Wide-ranged | Loss >50% | 1000 | 0.770 | 2100 | ssp585 |
| Woody | Wide-ranged | Loss <50% | 1000 | 0.182 | 2100 | ssp585 |
| Woody | Wide-ranged | Gain <50% | 1000 | 0.044 | 2100 | ssp585 |
| Woody | Wide-ranged | Gain >50% | 1000 | 0.004 | 2100 | ssp585 |

**Table S2. Proportion of Caatinga plant species showing projected range shift across different categories of woodiness and ecological generalism**. SSP denotes the future scenarios of Shared Socioeconomic Pathways. Range shift calculations considered the Caatinga extent.

| Woodiness | Generalism | Range shift Category | Number of Cells | % Species | Year | SSP |
|---|---|---|---|---|---|---|
| Non-woody | Narrow-ranged | Loss >50% | 251 | 0.761 | 2060 | ssp245 |
| Non-woody | Narrow-ranged | Loss <50% | 251 | 0.223 | 2060 | ssp245 |
| Non-woody | Narrow-ranged | Gain <50% | 251 | 0.016 | 2060 | ssp245 |
| Non-woody | Wide-ranged | Loss >50% | 884 | 0.191 | 2060 | ssp245 |
| Non-woody | Wide-ranged | Loss <50% | 884 | 0.594 | 2060 | ssp245 |
| Non-woody | Wide-ranged | Gain <50% | 884 | 0.212 | 2060 | ssp245 |
| Non-woody | Wide-ranged | Gain >50% | 884 | 0.003 | 2060 | ssp245 |
| Woody | Narrow-ranged | Loss >50% | 341 | 0.742 | 2060 | ssp245 |
| Woody | Narrow-ranged | Loss <50% | 341 | 0.226 | 2060 | ssp245 |
| Woody | Narrow-ranged | Gain <50% | 341 | 0.032 | 2060 | ssp245 |
| Woody | Wide-ranged | Loss >50% | 1000 | 0.290 | 2060 | ssp245 |
| Woody | Wide-ranged | Loss <50% | 1000 | 0.602 | 2060 | ssp245 |
| Woody | Wide-ranged | Gain <50% | 1000 | 0.106 | 2060 | ssp245 |
| Woody | Wide-ranged | Gain >50% | 1000 | 0.002 | 2060 | ssp245 |
| Non-woody | Narrow-ranged | Loss >50% | 251 | 0.861 | 2060 | ssp585 |
| Non-woody | Narrow-ranged | Loss <50% | 251 | 0.127 | 2060 | ssp585 |
| Non-woody | Narrow-ranged | Gain <50% | 251 | 0.012 | 2060 | ssp585 |
| Non-woody | Wide-ranged | Loss >50% | 884 | 0.279 | 2060 | ssp585 |
| Non-woody | Wide-ranged | Loss <50% | 884 | 0.515 | 2060 | ssp585 |
| Non-woody | Wide-ranged | Gain <50% | 884 | 0.201 | 2060 | ssp585 |
| Non-woody | Wide-ranged | Gain >50% | 884 | 0.005 | 2060 | ssp585 |
| Woody | Narrow-ranged | Loss >50% | 341 | 0.804 | 2060 | ssp585 |
| Woody | Narrow-ranged | Loss <50% | 341 | 0.173 | 2060 | ssp585 |
| Woody | Narrow-ranged | Gain <50% | 341 | 0.023 | 2060 | ssp585 |
| Woody | Wide-ranged | Loss >50% | 1000 | 0.395 | 2060 | ssp585 |
| Woody | Wide-ranged | Loss <50% | 1000 | 0.507 | 2060 | ssp585 |
| Woody | Wide-ranged | Gain <50% | 1000 | 0.094 | 2060 | ssp585 |
| Woody | Wide-ranged | Gain >50% | 1000 | 0.004 | 2060 | ssp585 |
| Non-woody | Narrow-ranged | Loss >50% | 251 | 0.884 | 2100 | ssp245 |
| Non-woody | Narrow-ranged | Loss <50% | 251 | 0.108 | 2100 | ssp245 |
| Non-woody | Narrow-ranged | Gain <50% | 251 | 0.008 | 2100 | ssp245 |
| Non-woody | Wide-ranged | Loss >50% | 884 | 0.329 | 2100 | ssp245 |
| Non-woody | Wide-ranged | Loss <50% | 884 | 0.486 | 2100 | ssp245 |
| Non-woody | Wide-ranged | Gain <50% | 884 | 0.180 | 2100 | ssp245 |
| Non-woody | Wide-ranged | Gain >50% | 884 | 0.005 | 2100 | ssp245 |
| Woody | Narrow-ranged | Loss >50% | 341 | 0.845 | 2100 | ssp245 |

| Woodiness | Generalism | Range shift Category | Number of Cells | % Species | Year | SSP |
|---|---|---|---|---|---|---|
| Woody | Narrow-ranged | Loss <50% | 341 | 0.144 | 2100 | ssp245 |
| Woody | Narrow-ranged | Gain <50% | 341 | 0.012 | 2100 | ssp245 |
| Woody | Wide-ranged | Loss >50% | 1000 | 0.471 | 2100 | ssp245 |
| Woody | Wide-ranged | Loss <50% | 1000 | 0.443 | 2100 | ssp245 |
| Woody | Wide-ranged | Gain <50% | 1000 | 0.082 | 2100 | ssp245 |
| Woody | Wide-ranged | Gain >50% | 1000 | 0.004 | 2100 | ssp245 |
| Non-woody | Narrow-ranged | Loss >50% | 251 | 0.976 | 2100 | ssp585 |
| Non-woody | Narrow-ranged | Loss <50% | 251 | 0.024 | 2100 | ssp585 |
| Non-woody | Wide-ranged | Loss >50% | 884 | 0.661 | 2100 | ssp585 |
| Non-woody | Wide-ranged | Loss <50% | 884 | 0.234 | 2100 | ssp585 |
| Non-woody | Wide-ranged | Gain <50% | 884 | 0.093 | 2100 | ssp585 |
| Non-woody | Wide-ranged | Gain >50% | 884 | 0.012 | 2100 | ssp585 |
| Woody | Narrow-ranged | Loss >50% | 341 | 0.947 | 2100 | ssp585 |
| Woody | Narrow-ranged | Loss <50% | 341 | 0.047 | 2100 | ssp585 |
| Woody | Narrow-ranged | Gain <50% | 341 | 0.006 | 2100 | ssp585 |
| Woody | Wide-ranged | Loss >50% | 1000 | 0.813 | 2100 | ssp585 |
| Woody | Wide-ranged | Loss <50% | 1000 | 0.147 | 2100 | ssp585 |
| Woody | Wide-ranged | Gain <50% | 1000 | 0.036 | 2100 | ssp585 |
| Woody | Wide-ranged | Gain >50% | 1000 | 0.004 | 2100 | ssp585 |

**SUPPLEMENTARY FIGURES**

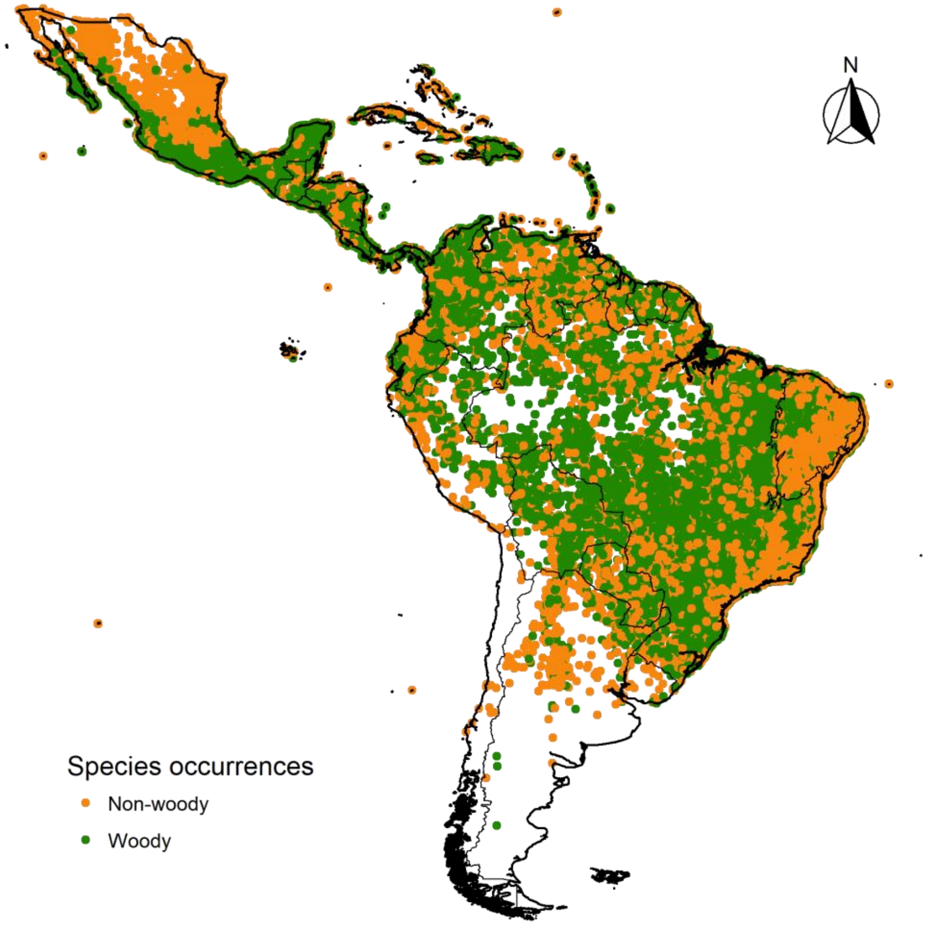


**Fig. S1. Species occurrence compiled for plant species known to occur within the Caatinga.** The map shows 335,091 records of 2,841 species categorized between two major growth-forms: Woody (trees, shrubs, palms, and woody vines) vs non-woody (herbs, herbaceous vines, and succulents) species.

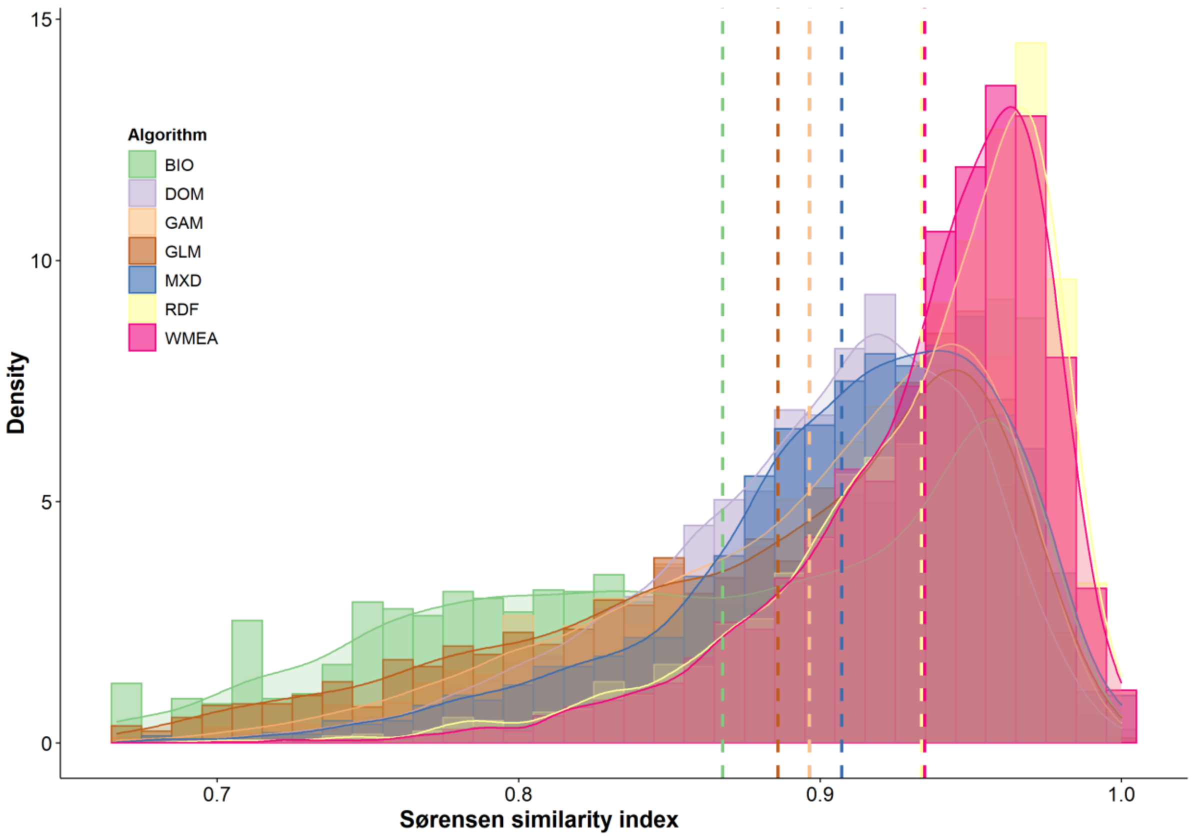


**Fig. S2. Performance metric of different algorithms across models for 2,841 plant species.** Vertical line denotes the mean value of Sørensen similarity index (performance metric) observed for the respective algorithm. Algorithm abbreviations: BIO = Bioclim climate envelope, DOM = Gower Environmental Distance - Domain, GAM = Generalised Additive Models, GLM = Generalised Linear Models, MXD = Maximum Entropy, RDF = Random Forests. WMEA represent the ensemble model computed as the weighted average across all six algorithms using the Sørensen similarity index as weight.

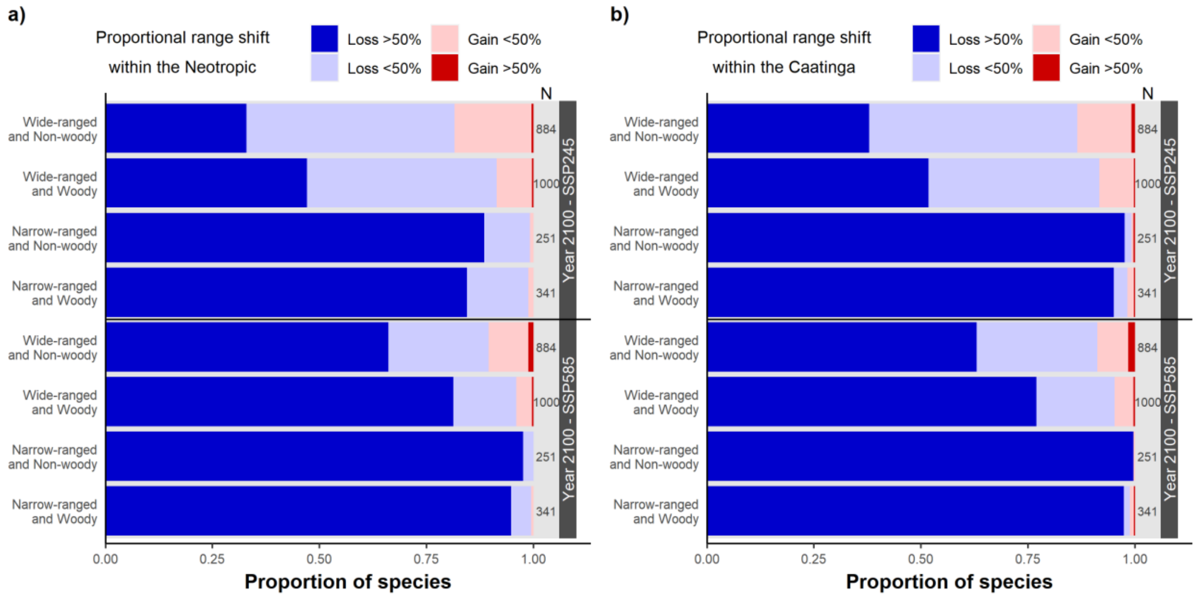


**Fig. S3. Projected range shift for species holding different levels of woodiness and ecological generalism.** Range shifts were computed separately within the Neotropics (a) and Caatinga (b). The results are shown for 2100 under the business-as-usual (SSP245) and non-mitigation (SSP585) scenarios. See Tables S1 and S2 for numeric details.

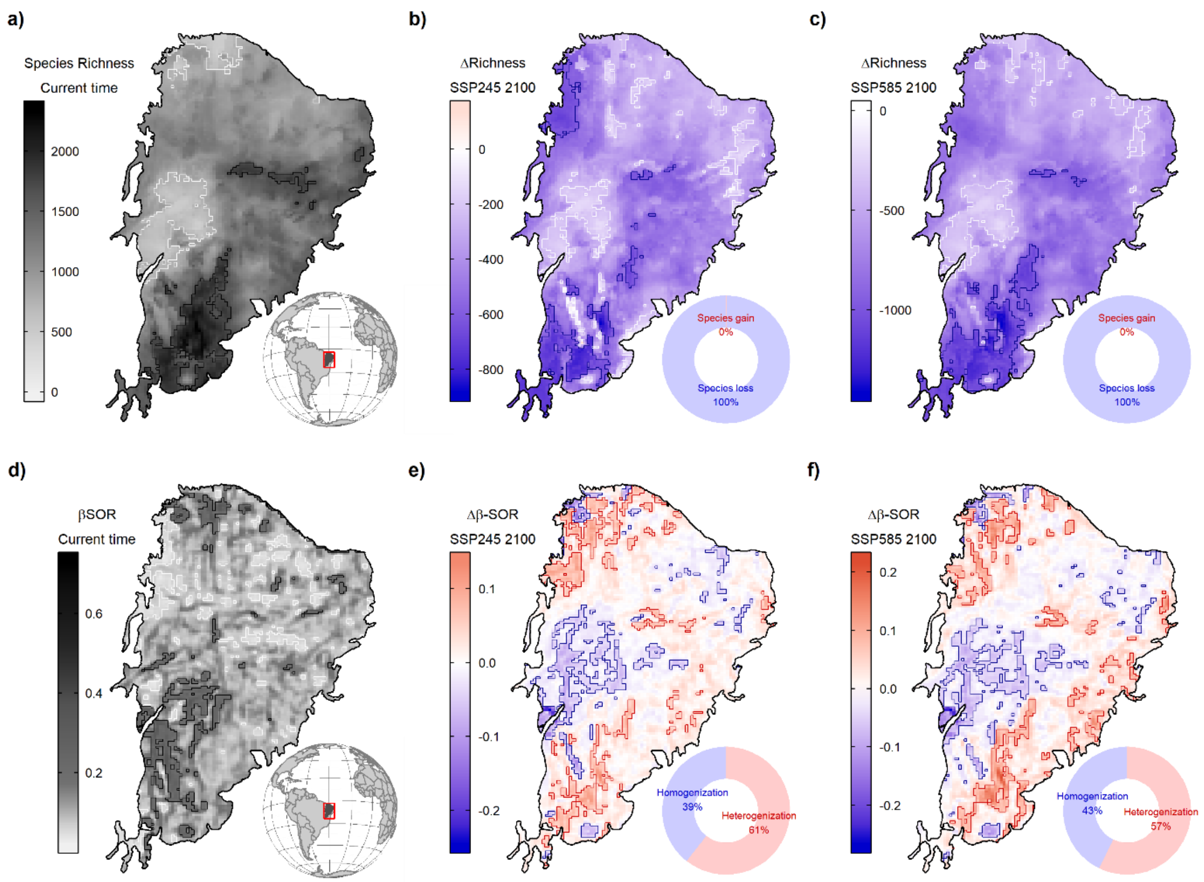


**Fig. S4. Geographical patterns of plant species richness and spatial-beta diversity in the Caatinga.** (a) Projected species richness at the current time. Expected change in species richness (ΔS) across plant assemblages under the (b) business-as-usual (SSP245) and (c) non-mitigation (SSP585) scenarios in 2100. (d) Spatial beta-diversity (βSOR) for the current time. Expected change in spatial beta-diversity (ΔβSOR) across plant assemblages under (e) SSP245 and (f) SSP585 scenarios for 2100. The contour lines denote the assemblages (cells) in the upper and lower 10% of the mapped pattern.

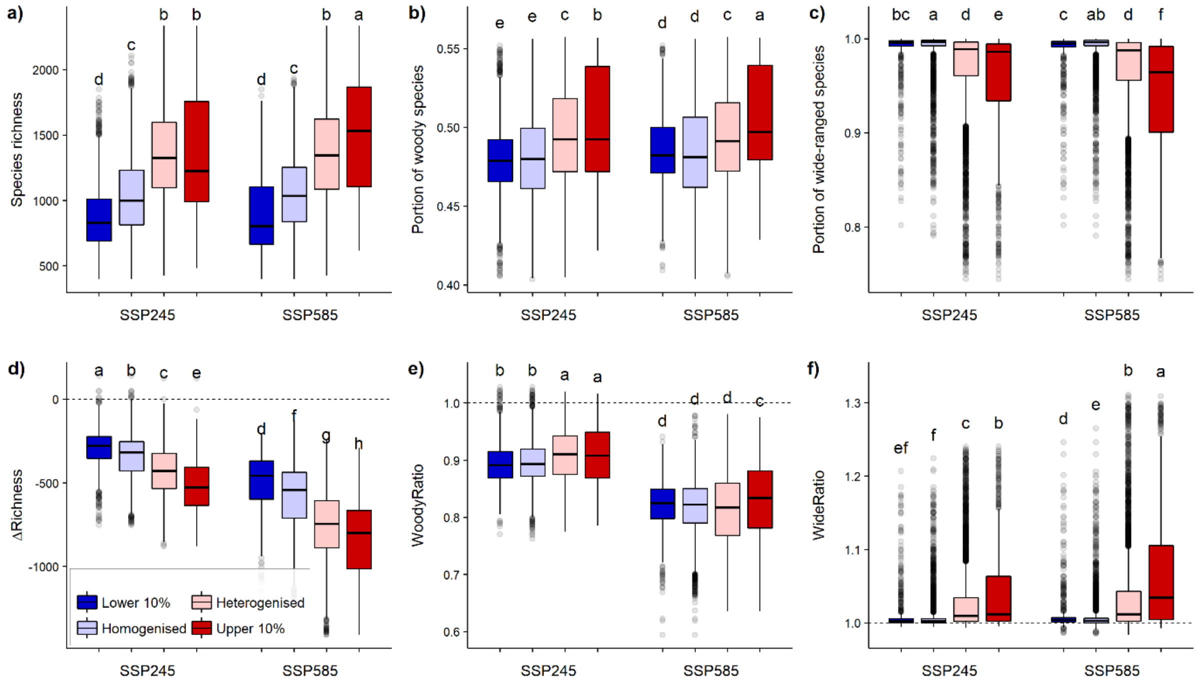


**Fig. S5. Assemblage-level metrics across regions subject to different levels of projected biotic change by 2100**. Each box denotes the median (horizontal line) and the 25th and 75th percentiles. Each box denotes the median (horizontal line) and the 25th and 75th percentiles. Vertical lines represent the 95% confidence intervals, and black dots are outliers. Small capital letters denote the results of the Kruskal–Wallis tests for the difference in medians across assemblages subject to different levels of biotic homogenisation: a) Species richness, $\chi^2$ = 4730.2, df = 7, p < 0.001. b) WoodyProp, $\chi^2$ = 1043.9, df = 7, p < 0.001. c) WideProp, $\chi^2$ = 3359.8, df = 7, p < 0.001. d) Δ Richness, $\chi^2$ = 10064.6, df = 7, p < 0.001. e) WoodyRatio, $\chi^2$ = 9622.8, df = 7, p < 0.001. f) WideRatio, $\chi^2$ = 2693.1, df = 7, p < 0.001. Woody and wide ratios above 1 indicate an increase in the assemblage-level proportion of woody and wide-range species in the future. All Kruskal–Wallis tests were subjected to Bonferroni correction of p-values.

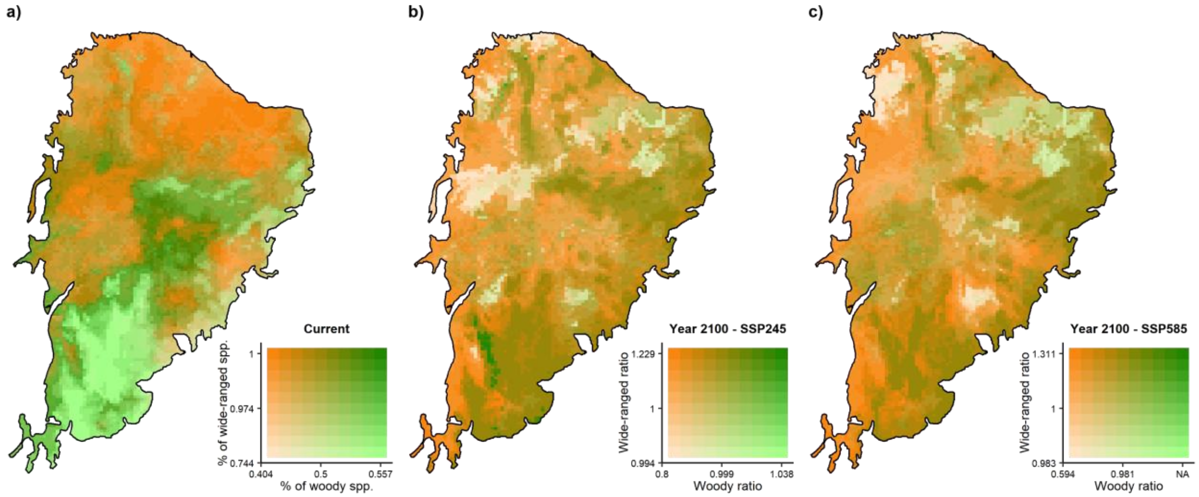


**Fig. S6. Patterns of woodiness and ecological generalism of plant assemblages in the Caatinga.** (a) Proportion of woody and wide-ranged species in plant assemblages. Relative change in the proportion of woody and wide-range species between 2100 and the current time under the (b) business-as-usual, SSP245 and (c) non-mitigation, SSP585 scenarios. Woody and Wide-range ratios above 1 indicate an increase in the assemblage-level proportion of woody and wide-range species in the future.

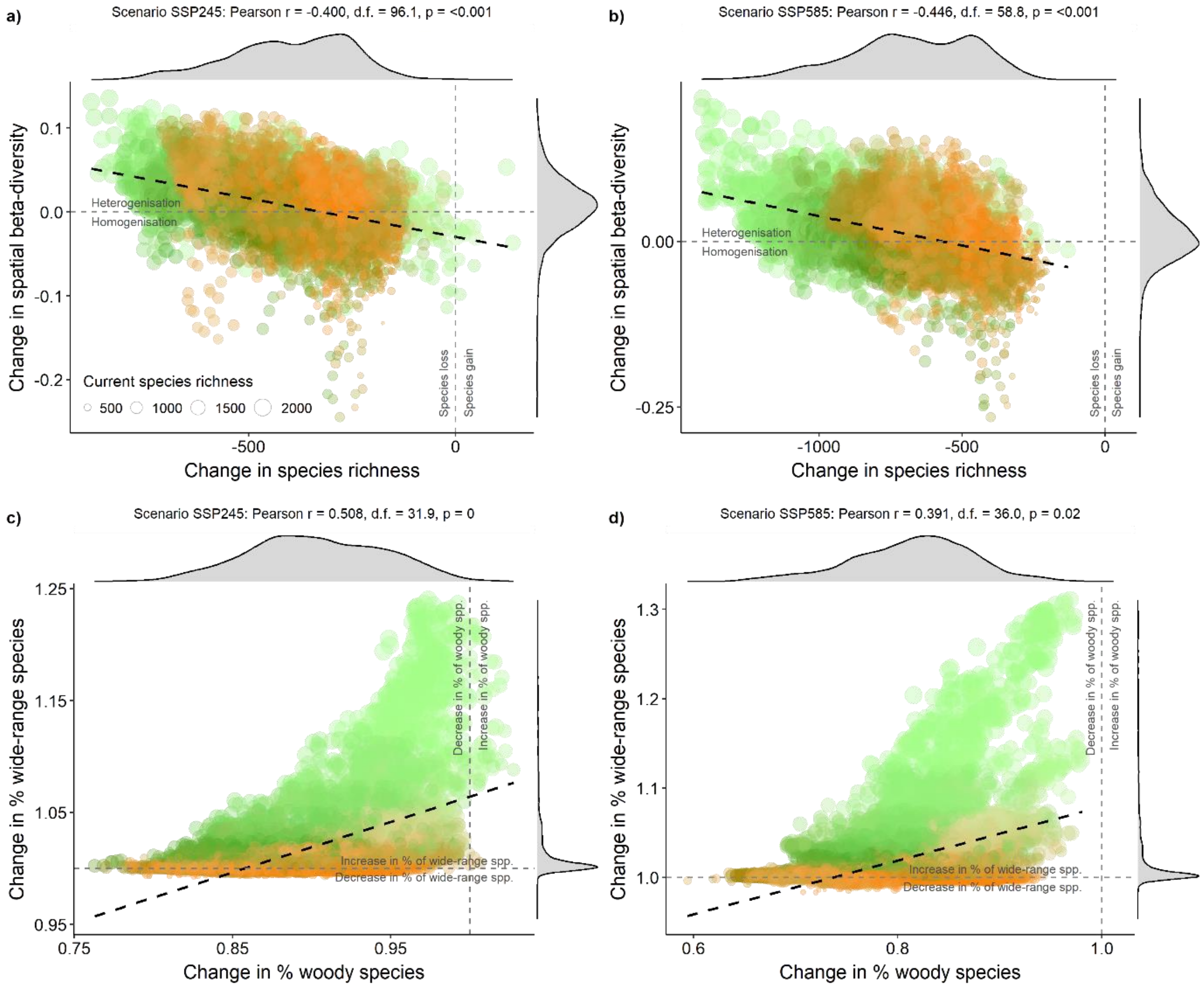


**Fig. S7. Projected change in species richness, spatial beta-diversity, woodiness, and ecological generalism in Caatinga plant assemblages by 2100.** Relationship between differences in species richness (ΔS) and spatial beta-diversity ($\Delta\beta_{SOR}$) in the scenarios (a) Business-as-usual [SSP245], and (b) Non-mitigation [SSP585]. Relationship between change in relative contribution of woody (WoodRatio) and wide-range (WideRatio) species. Symbol colours follow species assemblage representation in Fig. S6a.
